# Chloroplast genome analysis of Angiosperms and phylogenetic relationships among Lamiaceae members with particular reference to teak (*Tectona grandis* L.f)

**DOI:** 10.1101/2020.05.05.078212

**Authors:** P. Maheswari, C. Kunhikannan, R. Yasodha

**Affiliations:** Institute of Forest Genetics and Tree Breeding, Coimbatore 641 002 INDIA

**Keywords:** Phylogeny, chloroplast genome, angiosperms, *Tectona grandis*, Lamiaceae, nucleotide diversity

## Abstract

Availability of comprehensive phylogenetic tree for flowering plants which includes many of the economically important crops and trees is one of the essential requirements of plant biologists for diverse applications. It is the first study on the use of chloroplast genome of 3265 Angiosperm taxa to identify evolutionary relationships among the plant species. Sixty genes from chloroplast genome was concatenated and utilized to generate the phylogenetic tree. Overall the phylogeny was in correspondence with Angiosperm Phylogeny Group (APG) IV classification with very few taxa occupying incongruous position either due to ambiguous taxonomy or incorrect identification. Simple sequence repeats (SSRs) were identified from almost all the taxa indicating the possibility of their use in various genetic analyses. Large proportion (95.6%) of A/T mononucleotide was recorded while the di, tri, tetra, penta and hexanucleotide amounted to less than 5%. Ambiguity of the taxonomic status of *Tectona grandis* L.f was assessed by comparing the chloroplast genome with closely related Lamiaceae members through nucleotide diversity and contraction an expansion of inverted repeat regions. Although the gene content was highly conserved, structural changes in the genome was evident. Phylogenetic analysis suggested that *Tectona* could qualify for a subfamily Tectonoideae. Nucleotide diversity in intergenic and genic sequences revealed prominent hyper-variable regions such as, *rps16-trnQ*, *atpH-atpI*, *psc4-psbJ*, *ndhF*, *rpl32* and *ycf1* which have high potential in DNA barcoding applications.

---

Chloroplasts are the sunlight driven energy factories sustaining life on earth by generating carbohydrate and oxygen. Besides photosynthesis, chloroplast performs many biosynthesis such as fatty acids, amino acids, phytohormones, metabolites and production of nitrogen source (Daniell et al. 2016). Retrograde signalling of chloroplasts plays a significant role in biotic and abiotic stress responses in plants. Circular chloroplast (cp) genome with size range of 120 to 160 kb undergoes no recombination and uniparently inherited in angiosperms. It has been targeted for complete sequencing due to the importance of its gene content and conserved nature. The major features of angiosperm chloroplast genome include quadripartite circular structure with two copies of inverted repeat (IR) regions that are separated by a large single copy (LSC) region and a small single copy (SSC) region (Jansen et al. 2005). The genome includes SSRs, single nucleotide polymorphisms (SNPs), insertions and deletions (InDels), small inversions and divergent hotspots. The cp genome with GC content of 35 – 38 % encodes for 120-130 genes including protein coding genes, tRNA and rRNA genes. Genes with introns are 10-25 in number and few duplicated genes are also observed (Wicke et al. 2011). Knowledge on cp genome continues to reveal many variations and add information on their functional and evolutionary significance.

Sequencing of genome with unparalleled efficiency and precision provides enormous options to completely sequence cp genome. Several *de novo* assemblies of cp genome are reported frequently for many genome sequence deficient species thus providing a novel resource for functional genomics, evolutionary analysis and molecular breeding (Daniell et al. 2016; Guyeux et al. 2019). Unique information in cp genome allow overwhelming applications in taxonomy, phylogeny, phylogeography and DNA barcoding (Byrne and Hankinson 2012; Dong et al. 2012; Yan et al. 2019). Although the gene content and gene ordering in cp genome of plants is highly conserved (Daniell et al. 2016), several rearrangements occurred during evolution of plants (Ali 2019). In few plant families such as Fabaceae and Geraniaceae, the loss of one IR or few genes were observed, indicating the atypical evolution and gene functional changes among them (Martin et al. 2014). Further, accelerated mutation rate in certain regions of cp genome was also recorded. Both genic and intergenic regions show single nucleotide polymorphism (SNP), indels and simple sequence repeat (SSR) variations across and within plant species (Wang et al. 2018; Zhang et al. 2018a). SSRs are small repeating units of DNA harbouring high level variations in the sequence and used for species identification in many crop species (Shukla et al. 2018; Takahashi et al. 2018; Lu et al. 2018). Comparison of cp genome of two Apiaceae members, *Angelica polymorpha* Maxim. and *Ligusticum officinale* Koch showed the presence of a 418 bp deletion in the *ycf4*-*cemA* intergenic region of *A. polymorpha*, which was used in discrimination of these taxa (Park et al. 2019).

Most of the studies on cp genome are limited to solve specific problems at genera or family level. High resolution plant phylogenies have been reported to identify the relationship between the wild and cultivated taxa of economically important species (Carbonell-Caballero et al. 2015). Chloroplast genetic engineering has been proven as one of the successful options in crop genome modifications (Oldenburg and Bendich 2015; Daniell et al. 2016). Further, hybridization compatibility towards generation new cultivars can be assessed by their phylogenetic relationships. Geographical origin and history of domestication of a crop variety can be traced using cp genome which paves way for conservation and utilization of unique genetic variations (Wang et al. 2019; Nock et al. 2019). Recently, phylogeny of green plants were analysed with 1879 taxa (Gitzendanner et al. 2018) and 3654 taxa including the members of Chlorophyta, Charophyta, Rhodopyta, Bryophyta, Pteridophyta, Gymnospermae and Angiospermae (Yang et al. 2019). In the present study, cp genome sequences of 3265 Angiosperm taxa was analysed for their phylogenic relationships and distribution of simple sequence repeats (SSRs). Further, an attempt was made to elucidate the cp genome of *Tectona grandis* (teak), an economically important timber tree species growing in 60 different tropical countries, within the family Lamiaceae through phylogenetics, characterization of IR expansion and contraction and identification of genetic hotspot regions.

## Materials and Methods

The complete chloroplast genome sequences that belong to Magnoliophyta or Angiospermae were downloaded from NCBI on July 03, 2019 using the link https://www.ncbi.nlm.nih.gov/genome/browse#!/organelles/magnoliophyta. Two species of Gymnospermae *Taxus baccata* L. and *Pinus sylvestris* L. were included as outgroups. The sequence sampling included all the major lineages of Magnoliophyta belonging to 57 orders, 233 families and 3376 species. However, 111 taxa had the problems of duplicate genome entries or lesser number of genes and hence removed from the analysis. A complete list of 3265 taxa with two out group taxa with GenBank accession numbers is available in the Supplementary Table 1. SSRs in plastome sequence were mined using MISA, a Perl script (http://pgrc.ipkgatersleben.de/misa/misa). MISA detect microsatellites in FASTA formatted nucleotide sequence and generates output along with statistical data in two separate files. The MISA definition of microsatellites was set by unit size (x) and minimum number of repeats (y): 1/10, 2/7, 3/5, 4/5, 5/5, 6/5 (x/y). The maximal number of interrupting base pairs in a compound microsatellite was set to 100.

### Sequence Extraction, Alignment and Phylogeny analysis

Chloroplast genome phylogeny construction was performed with 60 genes extracted using BioEdit v7.0.5 (Hall 2011). The dataset included genes present in all the families like genes encoding small-ribosomal proteins (*rps2, rps3, rps4, rps8, rps11, rps14, rps15, rps18*), large ribosomal proteins (*rpl14, rpl20, rpl22, rpl32, rpl33, rpl36*), DNA dependent RNA polymerase (*rpoA, rpoB*), photosynthesis and energy production (*psaA, psaB, psaC, psaI, psaJ, psbA, psbB, psbC, psbD, psbE, psbF, psbH, psbI, psbJ, psbK, psbL, psbM, psbN, psbT, psbZ, atpA, atpB, atpE, atpH, atpI, ndhB, ndhC, ndhD, ndhE, ndhF, ndhG, ndhH, ndhI, ndhJ, ndhK, rbcL, petA, petG, petL, petN*) and others (*ycf4, matK, ccsA, cemA*).

The extracted gene sequences were aligned using MAFFT v7.409 (Katoh and Standley 2013) with the settings FFT-NS-2 and executed the output as FASTA format. The aligned sequences were trimmed to equal lengths at the ends using BioEdit sequence alignment editor software version 7.0.5 and the concatenated with MEGA X version 10.0.5 (Kumar et al. 2018). Phylogenetic analysis was performed for 3,265 species and outgroups with taxa, family and order levels using maximum parsimony method in PAUP* version 4.0b10 (Swofford 2002). The most parsimonious trees were recovered with a heuristic search strategy employing tree bisection and reconnection (TBR) with 100 random addition sequence replications. In an attempt to infer the taxonomic position of *Tectona grandis*, a subset of data consisting of only Lamiaceae family members along with two out group members were subjected to phylogeny analysis. All the phylogenetic trees were viewed and labelled using Interactive Tree Of Life (iTOL) v4 (https://itol.embl.de/) (Letunic and Bork 2019).

Gene distribution in cp genome of 10 Lamiaceae species selected as close relatives of *Tectona* was compared and visualized using mVISTA software in Shuffle-LAGAN mode (Frazer et al. 2004) with the annotation of *Tectona grandis* as a reference. The same alignment was used to calculate the nucleotide variability values (π) within Lamiaceae plastomes. The sliding window analysis was performed in DnaSP 6.10 (Rozas et al. 2017) with step size of 200 bp and window length of 600 bp and nucleotide diversity, π values were plotted using excel. Expansion and contraction of the IR regions among the selected Lamiaceae members were investigated and plotted using IRscope (Amiryousefi et al. 2018).

## Results and Discussion

### Organization, gene content, and characteristics of the chloroplast genome

Chloroplast genome of 3,265 taxa belonging to 56 orders and 223 families were analysed to understand the diversity and phylogenetic relationships among them. *Taxus baccata* and *Pinus sylvestris* were included as out groups for phylogeny analysis. Maximum and minimum plastome size of 2,42,575 bp and 1,13,490 bp was observed in *Pelargonium transvaalense* Knuth and *Aegilops cylindrica* L. respectively with the average size of 1,53,043 bp. The GC content was ranging from 33.6 to 43.6 with the average of 37.6%. Average gene content was 131 with maximum in *Perilla* (266 genes) and minimum (86 genes) in *Halesia diptera* Ellis (Supplementary Table 1). Sixty genes were considered for phylogeny analysis with minimum, maximum and average sequence length of 119 bp, 5591bp and 1366 bp respectively (Supplementary Table 2). The alignment of the concatenated data showed a minimum and maximum size of 37,700 bp (*Fargesia denudate* Yi) and 54,545 bp (*Carpinus tientaiensis* W.C.Cheng.) respectively with mean nucleotide composition of *A* = 28.1%, *C* = 17.4%, *G* = 20.3% and *T* = 31.6% (Supplementary Table 3).

Although cp genome information of green plants was employed for phylogeny analysis, consolidated data on SSR distribution across different orders is not available in the published literature. The assembled chloroplast genomes of 3265 species were mined for the presence of SSRs and a total of 163940 SSRs from 3265 scaffolds were identified (Table 1). Overall 119 repeat types were detected with two types of mono-nucleotides (A/T and G/C), three types of di-nucleotides (AC/GT, AG/CT and AT/AT), 12 types of tri-nucleotides, 6 types of tetra-nucleotides, 31 types of penta-nucleotides and 65 types of hexa nucleotides (Supplementary Table 4). The major SSR types with more than 20 numbers were listed in the Table 2. Among the identified SSRs, the mono-nucleotide was the most abundant SSR type, accounting for 95.6% of the total SSR motifs, which was followed by di (2.93%), tri (0.97%), tetra (0.076%) penta (0.13%) and hexa (0.23%) nucleotide SSR motifs. There was a large proportion of mononucleotide while the rest amounted to less than 5%. Although Poales had more number of SSRs, Asterales has diverse types. A/T richness of the chloroplast genome is identified in many previous studies (Cheon et al. 2019). It was suggested that such high amount of mononucleotide repeats in chloroplast genome may contribute to heritable variations (Bi et al. 2018).

### Phylogeny of Angiosperms

Application of phylogenetic reconstruction methods, unequivocal availability of genomic data and computational algorithms are continue to resolve several unanswered taxonomical problems in plant species. In the present study, phylogenetic trees had congruence with the Angiosperm Phylogeny Group (APG) IV system of classification. Six orders namely, Crossosomatales, Picraminiales, Metteniusales, Vahliales, Escalloniales and Bruniales did not have any representative taxa. Some of the orders such as Acorales, Amborellales, Aquifoliales, Berberidopsidales, Buxales, Canellales, Celastrales, Ceratophyllales, Chloranthales, Commelinales, Garryales, Huerteales, Icacinales, Oxalidales, Pandanales, Paracryphiales, Petrosaviales, Trochodendrales and Zygophyllales had 1-4 taxa representation. Different levels of diversification of angiosperms were recently proposed (Soltis et al. 2019) and the results obtained in this study showed high level of correspondence with ordinal level phylogeny of APG IV (Supplementary Fig. 1). The early diverging angiosperms including ANA grade, Magnoliids and Chloranthales formed a distinguishing clusters. COM clade comprising Celestrales, Oxalidales, and Malphigiales and Zygophyllales under Fabids formed a separate cluster along with nitrogen fixing clade Cucurbitales, Fabales, Fagales and Rosales. Early diverging eudicots Ceratophyllales, Ranunculales, Proteales, Trochodenrales, Buxales and Dilleniales formed a specific cluster. Similarly, superrosids and superastrids formed clearly distinguishable clusters. The monocot families Arecales, Acorales, Alismatales, Asparagales, Commelinales, Dioscoreales, Liliales, Pandanales, Petrosaviales, Poales and Zingiberales could be identified individually.

All the families within 56 orders were assessed for their taxonomical position and were in correspondence with the APG IV system of classification (Supplementary Fig. 2). However, at taxa level, few species were placed discordantly in different families. Recently, complete chloroplast genomes of 26 Gentianales species were analysed for phylogenetic relationships and found that *Gynochthodes nanlingensis* (Y.Z.Ruan) Razafim. & B.Bremer was grouped under Apocynaceae (Zhou et al. 2018). The present results were also confirmed the grouping of *G. nanlingensis* under Apocynaceae instead of Rubiaceae. Similarly, the genus *Wightia* of Scrophulariaceae has attracted attention in various studies (Zhou et al. 2014; Xia et al. 2019) and in the present study *Wightia speciosissimais* (D. Don) Merr. was placed closed to Phrymaceae members (Supplementary Fig. 2). Similarly, the genus *Pedicularis* of the family Orobanchasceae was grouped in Lamiaceae needs further studies. Recent report on molecular phylogeny of Orobanchaceae could not clear the species complex (Li et al. 2019). *Cypripedium macranthos* Sw. of Orchidaceae was surrounded by Asparagaceae members and *Lagerstroemia villosa* L. belonging to the family Lythraceae was embedded in between Combretaceae, could be due to taxonomical misidentification (Kim et al. 2014). Similarly, the species *Rivinia humilis* L. of the family Petiveriaceae and *Monococcus echinophorus* L. of the family Phytolaccaceae were placed conflictingly which reflected the uncertainty of their taxonomical status (Lee et al. 2013). *Monococcus echinophorus* was considered as a member of the family Petiveriaceae (Walker 2018). The taxonomic position of *Chaetachme aristata* Planch was established in the family Cannabaceae (Yang et al. 2013; Zhang et al. 2018b) instead Ulmaceae, and in the species tree it was grouped along with Cannabaceae members. All such inconsistent placement of plant species demands generation of cp genome information in all the plant taxa to suitably assess their phylogenic status.

### Taxonomy of Tectona

Till date the taxonomical status of the species *Tectona grandis* under Lamiaceae remain poorly understood phylogenetically. Complex morphological features of *Tectona* such as actinomorphic 5–7-lobed calyx and corolla, enlarged and inflated persistent calyx, and four chambered stony endocarp with small central cavity between the chambers posed problem in clear cut classifications. Many previous studies on molecular phylogeny of *Tectona* showed its unique phylogenetic position in Lamiaceae (Wagstaff and Olmstead, 1997; Li et al. 2016; Yasodha et al. 2018). In congruence with earlier studies, dendrogram obtained in the present study with 60 genes in 39 members of Lamiaceae confirmed the distinctness of the genus *Tectona* (Supplementary Fig. 3). All the lower level clades of phylogeny had 100% bootstrap values except *Tectona*-*Premna* clade showed 99.3%, establishing its difference from *Premna*. It was also confirmed that *Tectona* showed proximity to Premnoideae, Ajugoideaea and Scutellarioideae (Li et al. 2016).

Based on the results of phylogeny, 10 closely related taxa including *T. grandis* were selected for further characterization of cp genome. IRscope analysis is usually recommended for comparative cp genome analysis at species level to infer the structural variations in the large single copy (LSC), small single copy (SSC) and inverted repeats (IRs) junctions (Thode and Lohmann, 2019). This study depicted the genetic architecture of ten Lamiaceae members in the vicinity of the sites connecting the IRs to LSC and SSC regions to provide deeper insights of these junctions. The cp genome sequence of *Tectona grandis* (NC_020098.1) was 1,53,953◻bp, comprising LSC spanning 85,318◻bp, SSC spanning 17,741◻bp, and two IR regions each of 25,447◻bp length. Among the closely related species of *Tectona*, the region between IRb and LSC was conserved as *rpl22*-*rps19*-*rpl2* and the gene *rps19* extended in IRb with 36 to 61◻bp (Supplementary Fig. 4). The gene *ndhF* was placed in junction between IRb and SSC (JSB). The gene *ndhF* was present in SSC region of *Tectona grandis* whereas in *Premna microphylla* it was present 6bp away from IRb region. The JSA junction (SSC/IRa) and JLA junction (LSC/IRa) was characterized by the presence of *ycf1* and *trnH* gene respectively, except *Tecona grandis*, where tRNA was present in JLA junction and loss of *trnH* gene was observed. In *Scutellaria baicalensis*, the *trnH* was present in LSC region. Duplication of the genes in cp genome is widely reported (Raveendar et al. 2015) and duplicated *rps19* and *ycf1* among the analysed species was obvious. Chloroplast genome evolution is governed by the IR expansions and contractions leading to variations in the genome size (Könyves et al. 2018; Li et al. 2018). However in this study, LSC region (81,770 bp to 86,078 bp) had wide variation, whereas less variation in IR and SSC regions were observed. Further within genus such as *Holcoglossum*, no variations in IR regions were reported (Li et al. 2019).

Level of sequence divergence among the cp genome would provide the basic information on similarity across individuals. Sequence divergence of *T. grandis* was assessed by the comparative analysis among the closely related Lamiaceae members using mVISTA. The cp genomes showed sequence divergence below 50%, indicating a low conservatism in the non coding regions of these chloroplast genomes (Fig. 1). The alignment revealed species sequence divergence across the cp genome, signifying the lack of conservation of genome. The single-copy regions, intergenic regions and genic regions were more divergent except for few genic regions like *rrn16*, *rrn23*, *ndhB* and, *rps7* (Fig. 1). According to the chloroplast genome sequence alignment of the ten Lamiaceae taxa, about 29 hyper-variable regions were discovered with nucleotide diversity (π) value of 0.07 and above (Fig. 2). Some of the important hotspot regions were *rps16-trnQ*, *atpH-atpI*, *psc4-psbJ*, *ndhF*, *rpl32* and *ycf1*. These mutational hotspots would serve as potential loci for developing novel DNA barcodes for plant classification. Thus, phylogeny and distinctness of the cp genome of *Tectona* confirmed the earlier proposal of a new subfamily Tectonoideae with *Tectona* as a monotypic genus (Li and Olmstead 2017).

**Fig 1.**
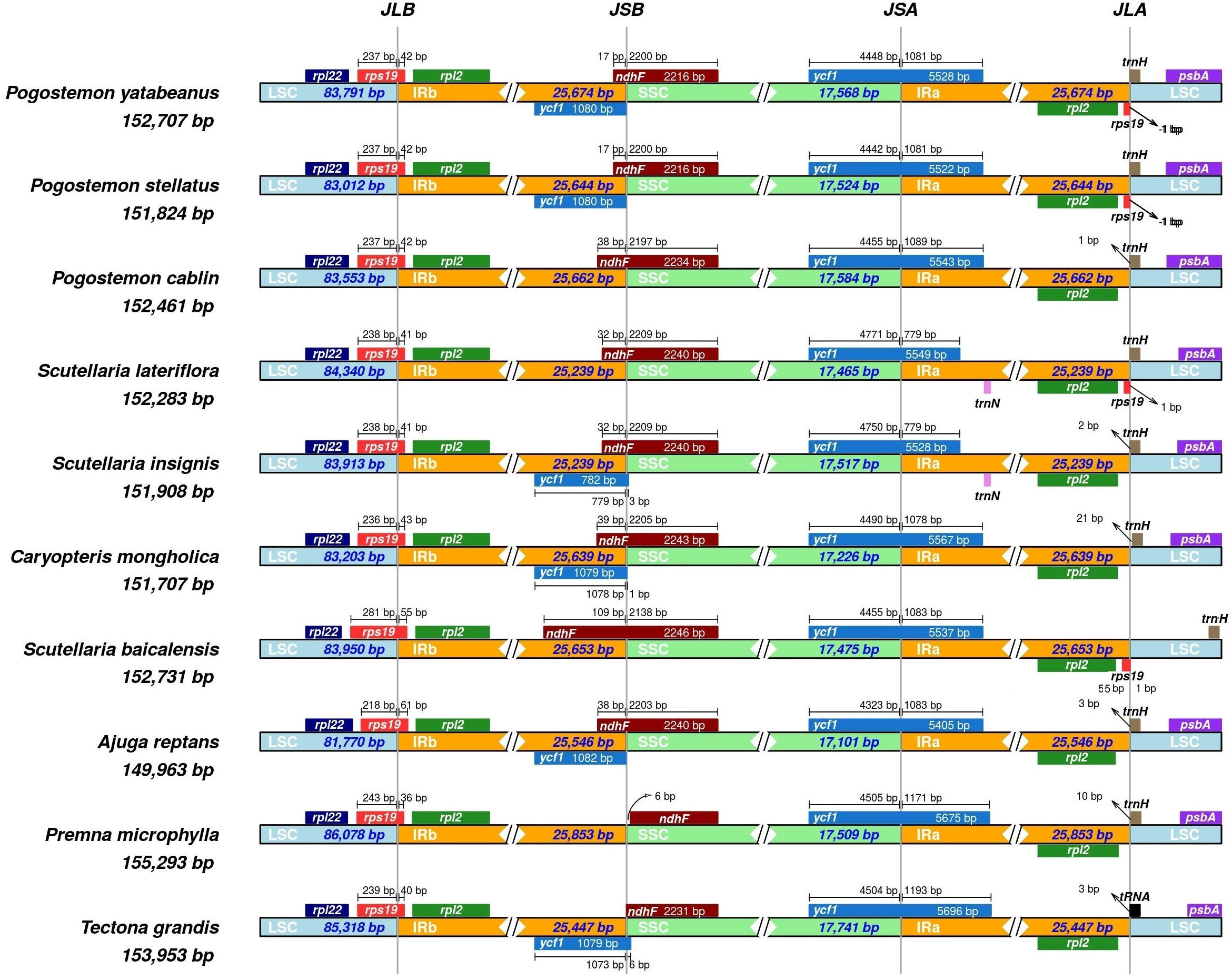
Comparisons of the Large Single Copy (LSC), Inverted Repeat a (IRa), Small Single Copy (SSC), and Inverted Repeat b (IRb) boundaries among ten Lamiaceae chloroplast genomes. Boxes above the main line represent the genes at the IR/SC borders.

**Fig. 2.**
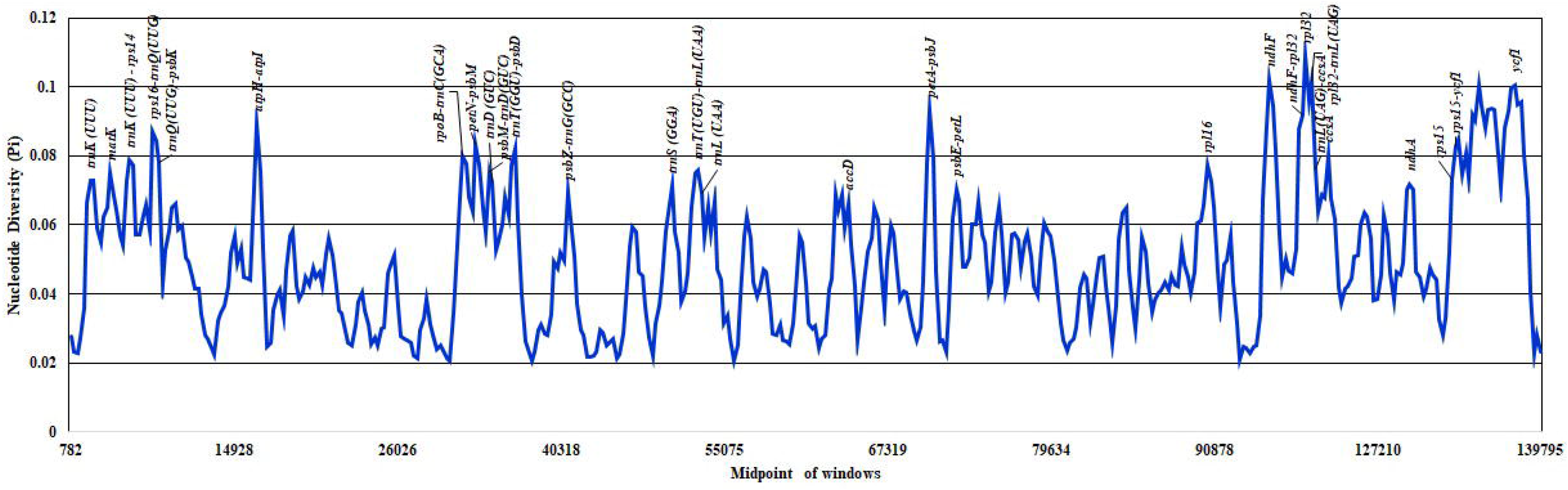
Sliding window analysis of the whole chloroplast genome of 10 species of Lamiaceae members. (window length: 600◻bp, step size: 200◻bp). X-axis: position of the midpoint of a window, Y-axis: nucleotide diversity of each window.

## Conclusion

Unprecedented availability of genomic data from organelles provides opportunities to discern the evolutionary relationships among the plant species. In the current study, chloroplast genome phylogeny of 3625 Angiosperm taxa revealed the phylogenetic and taxonomic status in congruence with APG IV classification. Valuable data generated on distribution of SSRs would give an impetus on their use in species identification and evolution of several angiosperm taxa. Many plant species requiring deeper understanding on phylogeny and taxonomy were identified and generation of more cp genome data in many of the unexplored plant species becomes essential. The phyogeny of Lamiaceae members, variations in structure and gene content confirmed the unique status of *Tectona* in the family. Nucleotide diversity among the closely related species in Lamiaceae with several variable hotspots would be useful to develop DNA markers suitable for discrimination of species and inference of phylogenetic relationships.

## Supporting information

Fifty percent maximum parsimony majority rule consensus tree of Angiospermae with major clades inferred from 60 chloroplast protein coding genes.

Phylogenetic tree of 3265 taxa data set with two outgroups based on 60 chloroplast protein coding genes using maximum parsimony (MP).

Identity plot comparing 10 Lamiaceae members chloroplast genome sequences with annotations, using mVISTA.

Phylogenetic tree of Lamiaceae members.

## Acknowledgements

The authors acknowledge the Department of Biotechnology (DBT), Government of India for financial support. Junior Research Fellowship provided to P. Maheswari by DBT is acknowledged.

## List of Supplementary Tables

Supplementary Table ESM_1: List of Angiospermic plant taxa used for phylogeny study

Supplementary Table ESM_2: List of genes in the chloroplast genome of the 3265 taxa

Supplementary Table ESM_3: Information on concatenated dataset for the 3265 taxa

Supplementary Table ESM_4: Details on simple sequence repeats present in the chloroplast genome analysed

## List of Supplementary Figures

**Supplementary Fig. 1. Fifty percent maximum parsimony majority rule consensus tree of Angiospermae with major clades inferred from 60 chloroplast protein coding genes**. Terminals with a circle represent collapsed clades with◻≥◻2 taxa.

**Supplementary Fig. 2. Phylogenetic tree of 3265 taxa data set with two outgroups based on 60 chloroplast protein coding genes using maximum parsimony (MP)**. The coloured strips indicate the clustering of the MP tree at the family and ordinal level. Ordinal and higher-level group names follow APG IV.

**Supplementary Fig. 3. Phylogenetic tree of Lamiaceae members.** Dendrogram generated based on whole chloroplast genome using maximum parsimony (MP) with 50% majority‐rule consensus principle.

**Supplementary Fig. 4. Identity plot comparing 10 Lamiaceae members chloroplast genome sequences with annotations, using mVISTA.** The vertical scale indicates the percentage of identity, ranging from 50 to 100%. The horizontal scale indicates the coordinates within the chloroplast genomes. Grey arrows represent the genes and their orientations. Blue boxes represent exon regions and red boxes represent non-coding sequence (CNS) regions.

