## Supplementary figures and images for "Chloroplast genome analysis of Angiosperms and phylogenetic relationships among Lamiaceae members with particular reference to teak (*Tectona grandis* L.f)"

### Fifty percent maximum parsimony majority rule consensus tree of Angiospermae with major clades inferred from 60 chloroplast protein coding genes.

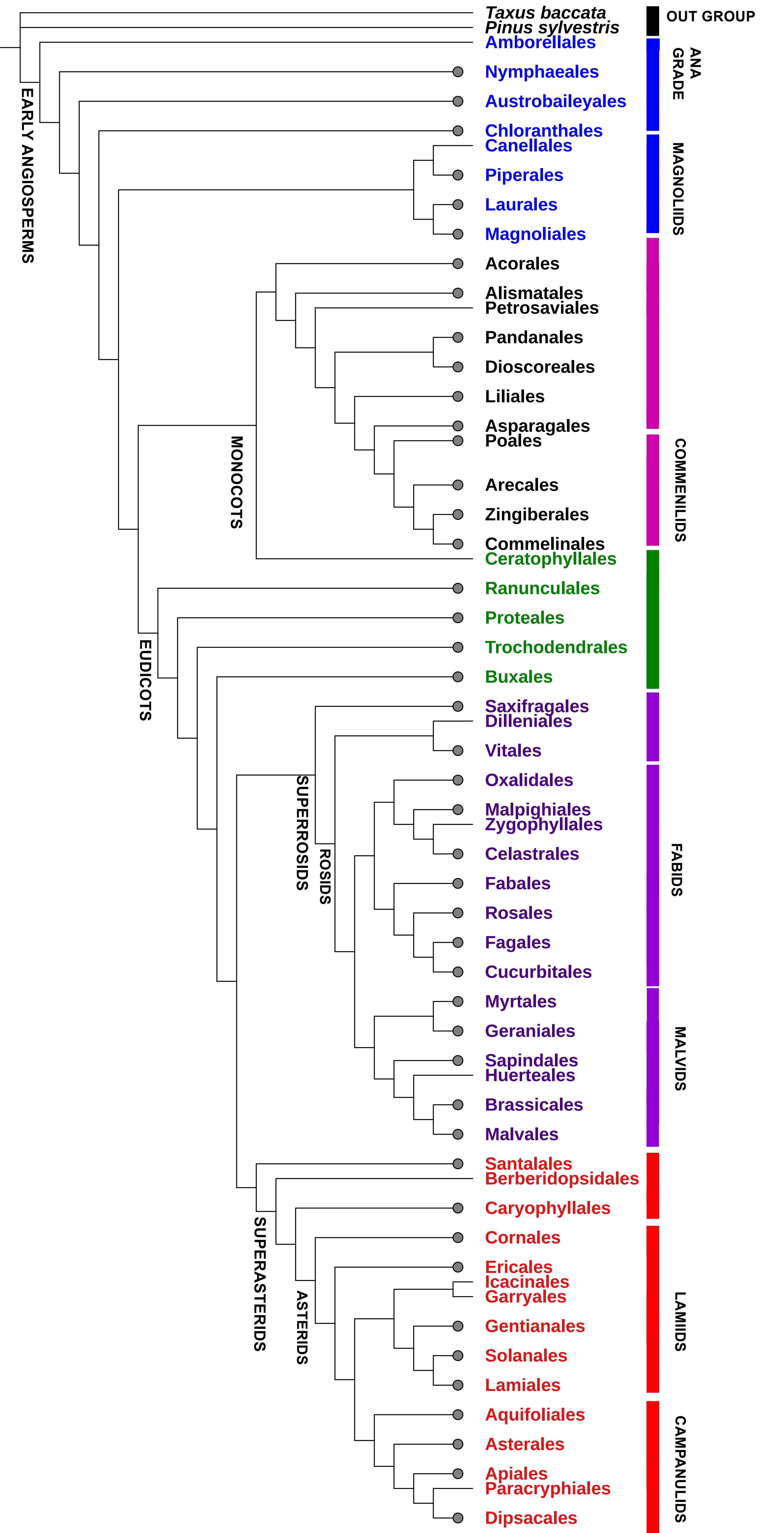

### Identity plot comparing 10 Lamiaceae members chloroplast genome sequences with annotations, using mVISTA.

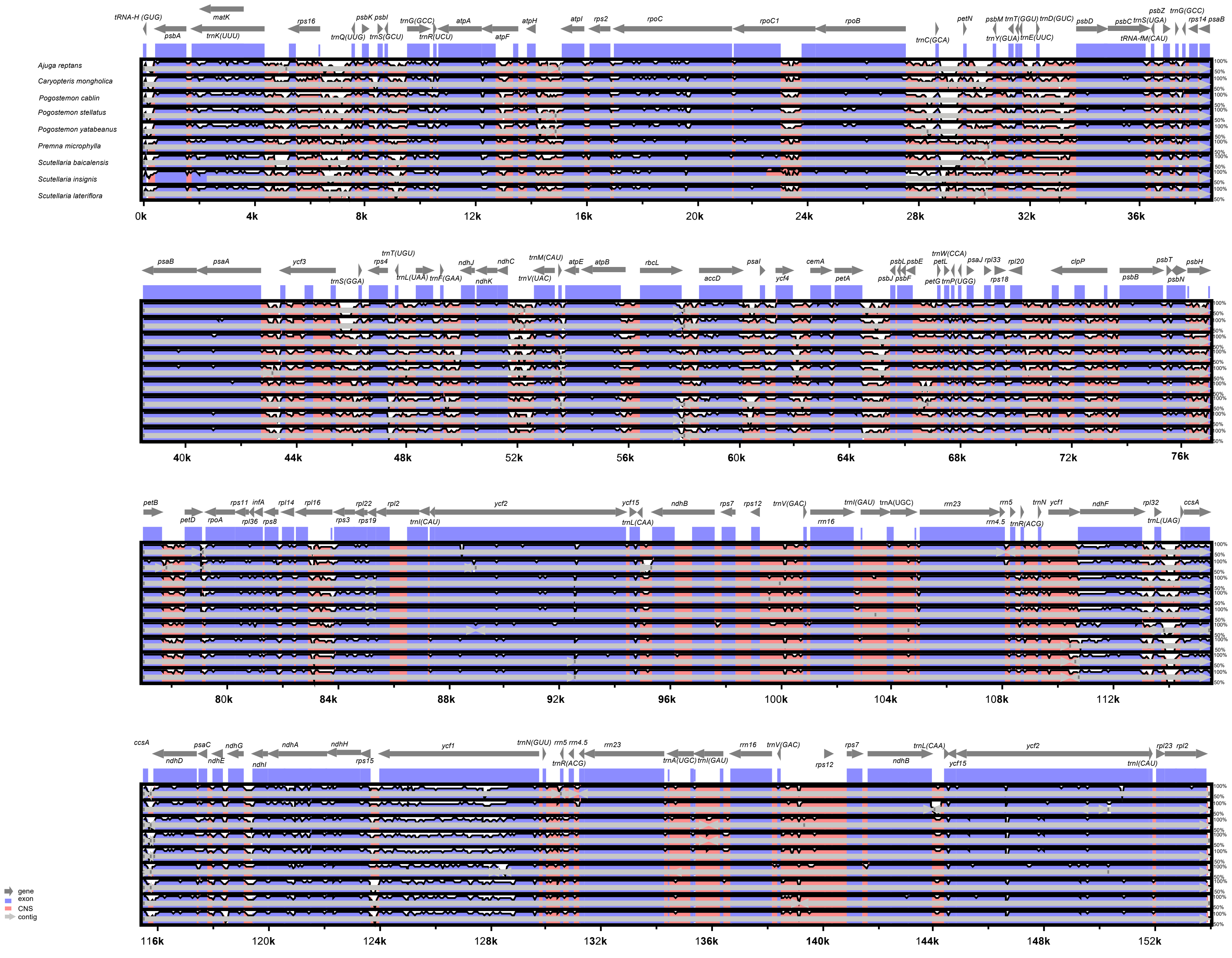

### Phylogenetic tree of 3265 taxa data set with two outgroups based on 60 chloroplast protein coding genes using maximum parsimony (MP).

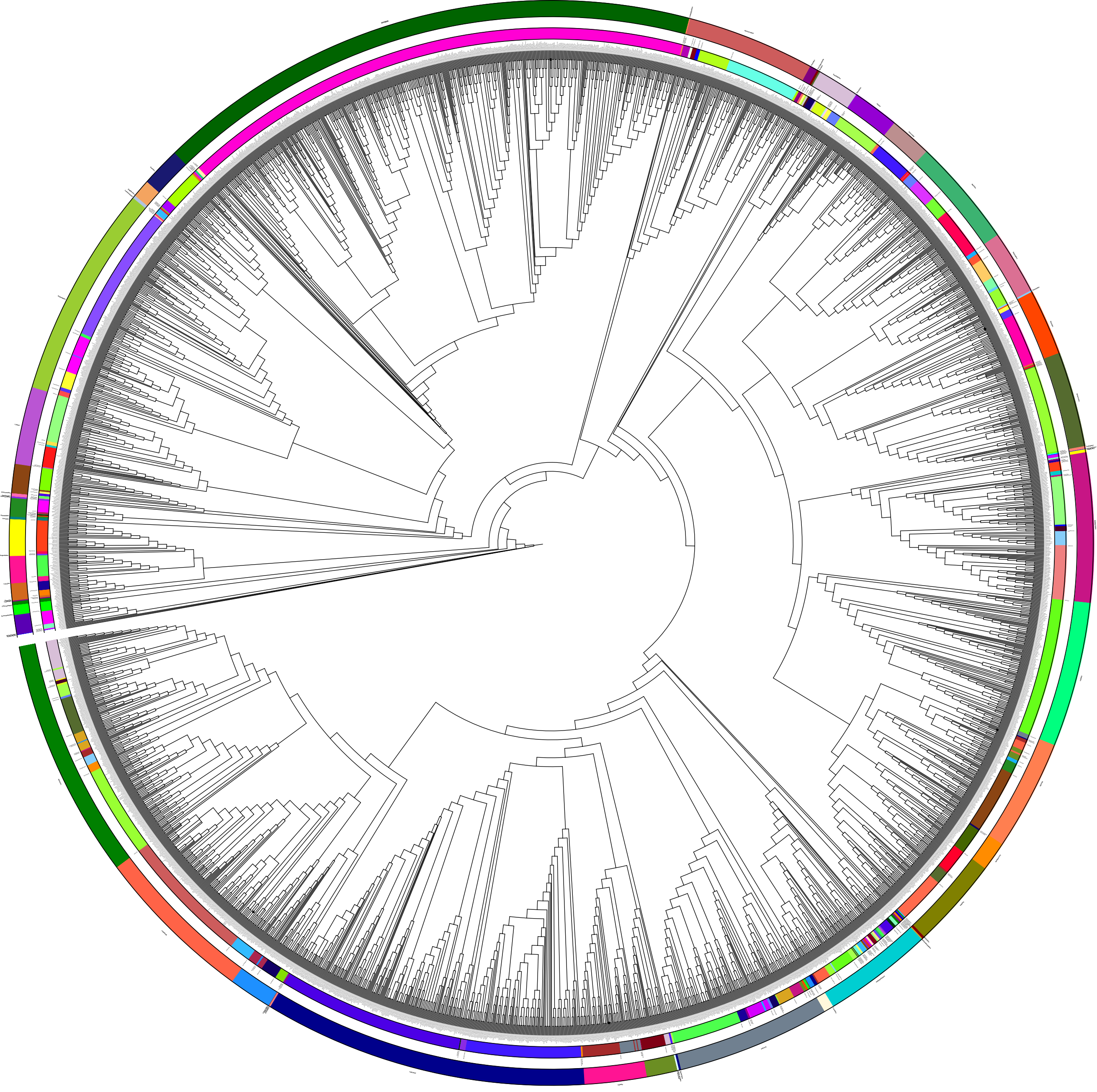

### Phylogenetic tree of Lamiaceae members.

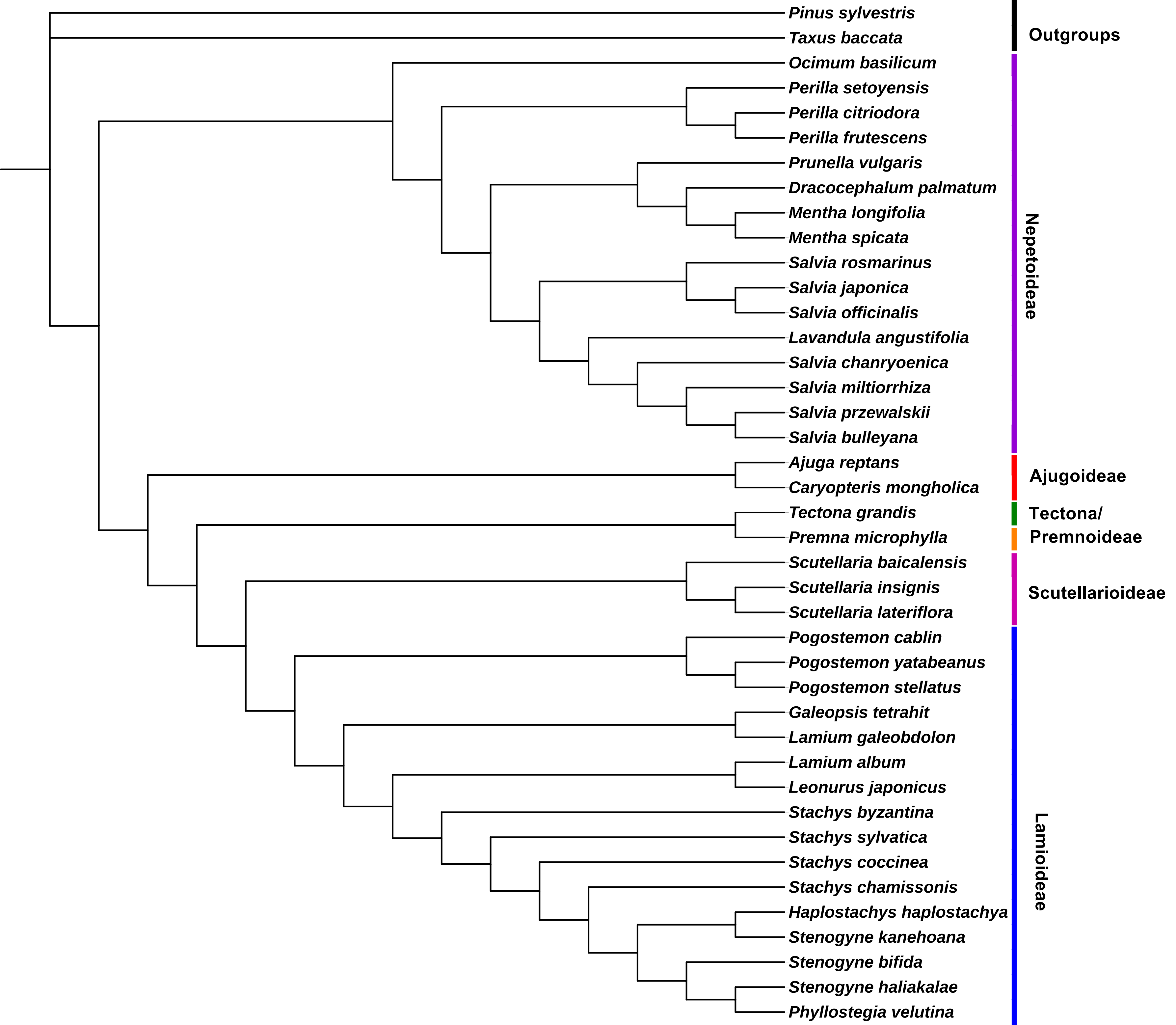
